# Medial septum glutamate neurons are essential for spatial goal-directed memory

**DOI:** 10.1101/2022.03.16.484657

**Authors:** Jean-Bastien Bott, Jennifer Robinson, Frederic Manseau, Etienne Gauthier-Lafrenière, Sylvain Williams

## Abstract

The medial septum (MS), a brain region containing acetylcholine, GABA and glutamate neurons, is essential for learning and memory. However, whether MS glutamate neurons contribute to memory is unknown. Here, we use calcium imaging and optogenetic silencing to determine the function of MS glutamate neurons in mice performing a spatial allocentric memory navigation task. While MS glutamate neurons appeared randomly active during free exploration, two groups of glutamate neurons emerged during training in a five-arm star maze, distinguished by their peak activity. The largest group was predominantly activated before locomotion whereas the other was primarily active when mice reached the reward site. Both populations were preferentially activated for correct over incorrect trajectories. Interestingly, optogenetic silencing of MS glutamate neurons immediately before mice start navigating, induced significant spatial memory impairments. Together these results demonstrate that MS glutamate neurons heterogeneously encode navigationally relevant information and are essential for spatial learning.

## INTRODUCTION

The medial septum (MS) is a major input pathway to the hippocampus network and has an essential role in learning and memory (Winson, 1978). The MS is composed of three different neuronal populations: GABAergic, cholinergic and glutamatergic neurons (Freund & Antal, 1988; Kimura et al., 1980; Sotty et al., 2003), with MS glutamate neurons making up 25% of the total septal population and accounting for approximately 1/5 of the projections from the septum to the hippocampus (Sotty et al., 2003). MS glutamate neurons have both local and long-range projections (Sotty et al., 2003; Manseau et al., 2005, Huh et al., 2010; Fuhrmann et al., 2015; Robinson et al., 2016) and can modulate hippocampal theta frequency rhythm (Fuhrmann et al., 2015, Robinson et al., 2016). Recent evidence suggests that MS glutamate neurons may promote locomotion, as shown by activity recorded at the population level with fiber-photometry in both head-fixed and freely-moving animals (Fuhrmann et al., 2015; Zhang et al., 2018) and may also provide speed correlated inputs to the medial entorhinal cortex (Justus et al., 2017). In addition, MS glutamate neurons have been shown to project to the lateral habenula and when activated, promote aversion behaviours (Zhang et al., 2018). However, it remains unknown if the MS glutamate neurons contribute to navigation and memory function. Here, to examine coding activity of MS glutamate neurons during free exploration and spatial learning, we performed calcium imaging with miniature microscopes and optogenetic manipulations during an episodic-like spatial learning and memory task in a 5-arm star-maze. The task was divided into three learning phases allowing to examine MS glutamate contribution to new spatial memory formation (initial learning), remote memory (retraining) and memory updating (reversal learning). Using single-cell resolution calcium imaging, we demonstrate that MS glutamate neurons consist of two main subpopulations with distinctive activation patterns across the learning phase of the task. While only a small population was primarily activated during locomotion, the largest populations of MS glutamate neurons was active predominantly with other periods of the task, specifically, before the onset of locomotion in the start-arm or at the end of locomotion when mice entered the destination arm. Interestingly, MS glutamate activity before locomotion onset or at the destination arm were higher depending if mice performed a correct trajectory toward the target arm. Finally, to determine if the activity of MS glutamate neurons preceding locomotion onset was a key contributor to task performance, this population was optogenetic silenced only while animals were in the start arm before each trial began. Remarkedly, silencing MS glutamate neurons during this period immediately preceding active navigation resulted in significant learning and memory deficits. Together these results demonstrate that there are two principal populations of MS glutamate neurons; *preparatory cells* that are preferentially active before correct trajectory onset and are essential for successful goal-directed spatial navigation and *target cells* active when mice reached the goal region.

## RESULT

### Ca^2+^ imaging of MS glutamate neurons in a goal-directed spatial memory task

MS glutamate neurons were transfected with the calcium indicator GCAMP6f using a Cre-dependent Adeno-associated virus in VGLUT2-Cre(KI)-mice (**Fig. 1a**; methods). The activity of MS glutamate neurons was visualized at single-cell resolution with head-mounted epifluorescent miniature microscopes in freely-behaving mice (**Fig. 1a and 2a**; methods). Mice were recorded during a new dry and allocentric-version of the Star-Maze task (Rondi-Reig et al., 2006; methods), where mice were trained to navigate towards a single target arm among five which contained a water reward. The task was organized in three successive learning phases (**Fig. 1a**): i) an initial learning (d1-4) during which mice have to learn how to navigate to the target arm; ii) a re-training (d29-32) during which mice are tested for long-term spatial memory; iii) a reversal learning (d33-36) allowing to measure mice ability to form a new spatial memory. Each phase consisted of 3 days of learning with 4 trials per day, followed by a probe trial on the fourth day (**Fig. 1a**; methods). For each trial, water-deprived mice were first held for 30s in a pseudo-randomly selected start arm. Following the 30s initial start period, the door opened, and mice were left to freely explore the 5-arm star maze until the target arm containing a water-reward was reached. On the fourth day (the probe trial), a similar experiment was done except the water reward was absent. (**Fig. 1b**; methods).

**Figure 1.**
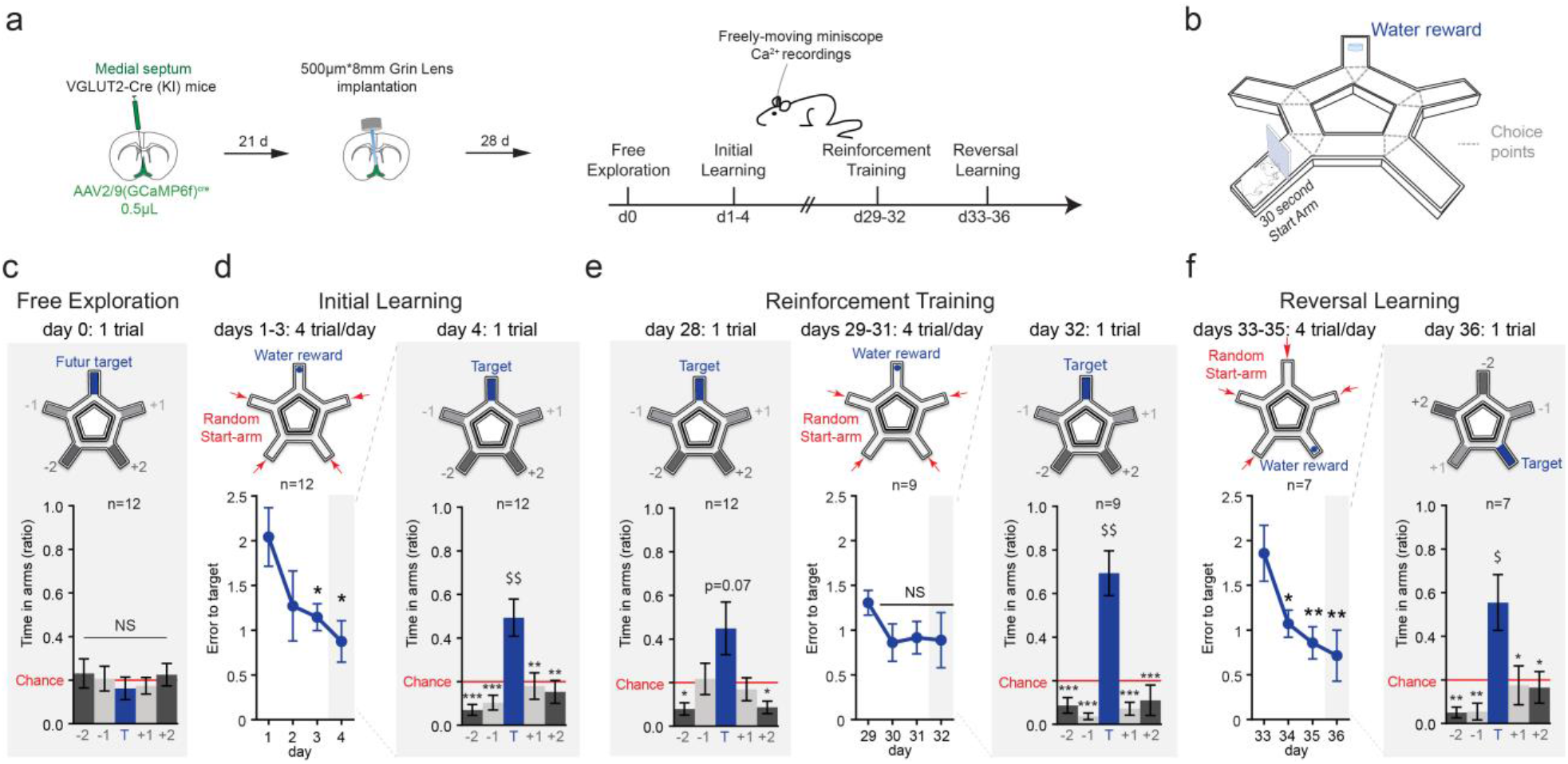
Freely-moving Ca^2+^ imaging of medial septum glutamatergic neurons during a spatial navigation learning task. **a**, Experimental time-line.Spatial memory testing was divided in three phases: i) initial learning, ii) retraining, iii) reversal learning. **b**, Trial began with a 30s waiting period in a pseudo-randomly selected Start-Arm. Then water-restricted mice freely-explore the maze to find the Target-Arm. **c**, Free exploration. Time spent in each arm during the free-exploration trial did not differ. **d**, Initial Learning. Left: across learning days, mice reduced the number of error before entering the target arm (*Holm-Sidak’s multiple comparison*). Right: During recent memory probe trial mice spent more time in target arm than in other (*Holm-Sidak’s multiple comparison*) or than chance (One-sample t-test; T_(11)_=3.435; p=0.0056). **e**, Re-training. Left: during the remote memory probe trial, mice still displayed a preference for the target arm compared to the other (*Holm-Sidak’s multiple comparison*) but only tend to visit it more than chance (One-sample t-test; T_(8)_=2.065; p=0.0728). Middle: Across re-training days, mice did not reduce further the number of error before entering the target arm (*Holm-Sidak’s multiple comparison*). Right: during the following memory probe trial mice displayed a strong preference for the target arm compared to the other (*Holm-Sidak’s multiple comparison*) and to chance level (One-sample t-test; T_(8)_=4.788; p=0.0014). **f**, Reversal learning. Left: across reversal learning days (shift of the target arm position), mice reduced the number of entries in error arms before entering the new target arm location (*Holm-Sidak’s multiple comparison*). Right: during the following memory probe trial mice displayed a preference for the new target arm location compared to the other (*Holm-Sidak’s multiple comparison*) and to chance level (One-sample t-test; T_(6)_=2.773; p=0.0323). Error bars represent s.e.m. *P<0.05; **P<0.01; ***P<0.001 (*Holm-Sidak’s multiple comparison*). $P<0.05; $$P<0.01 (One-sample t-test vs. chance).

**Figure 2.**
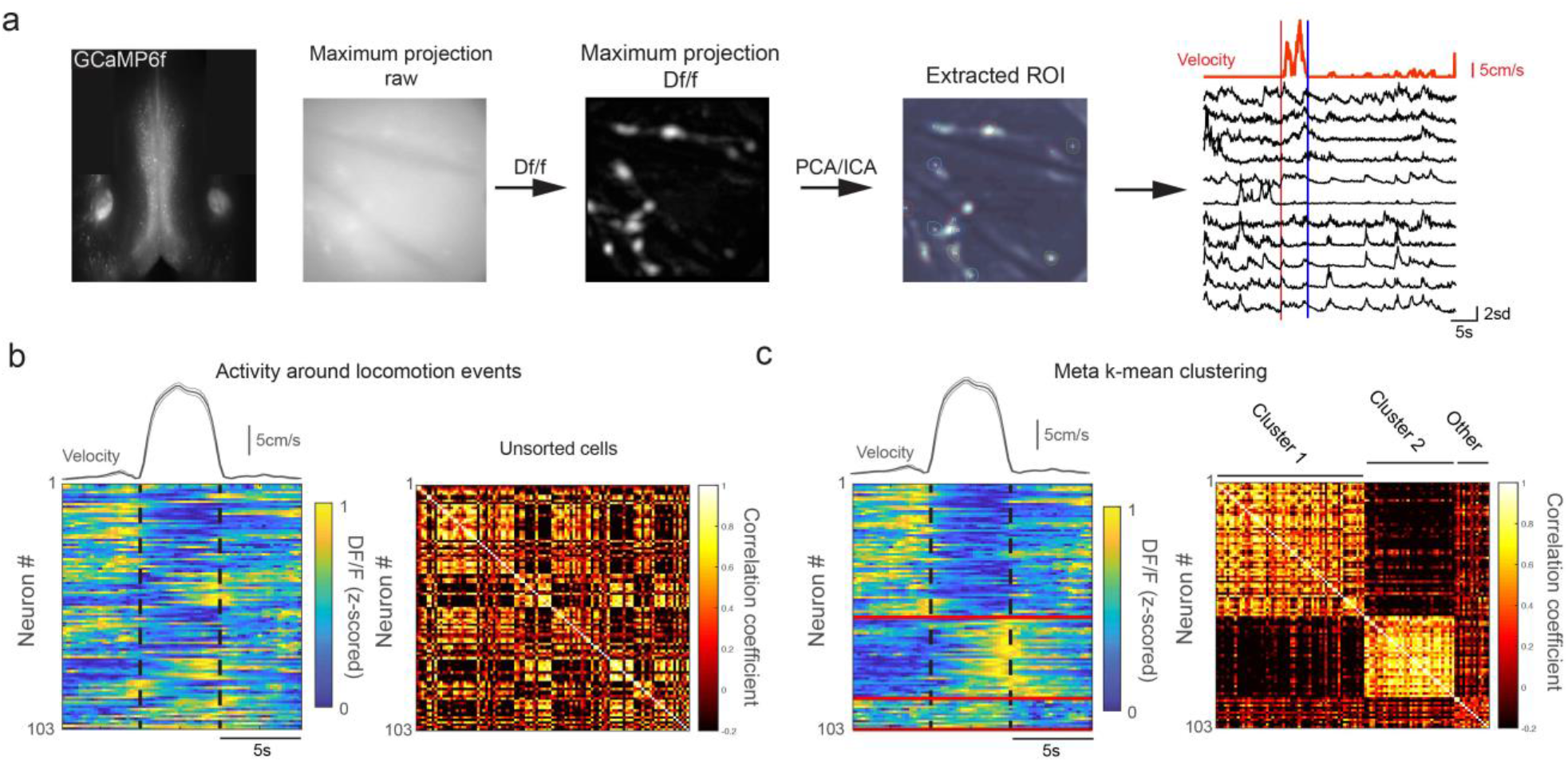
Functional subpopulations of MS glutamate neurons. **a**, transfected fluorescent medial septum area following GCaMP6f injection in VGlut2-Cre mice. Images recorded using a miniature microscope. Raw images were converted into DF/F movies and cells were automatically detected and segmented using an PCA/ICA algorithm. Right: Example cell activity from one mouse (in black) around a locomotion event (velocity in red) on day 2 of the initial learning phase. **b**, realigned mean activity around all locomotion events for all cells; left: each row represents the averaged Ca^2+^ response of a single neuron, with color representing fluorescence intensity (z-scored, 103 cells from 12 mice); right: neuron-neuron mean trace correlation. **c**, after meta-K-means clustering, neurons were grouped into 3 clusters based on their temporal activity. Neurons were sorted based on their respective cluster.

We first showed that naïve mice that were freely-exploring the device (d0) displayed no preference for the future target arm (**Fig. 1c**; *Arm effect*; F_(4;44)_=0.2592; p=0.9025). During the initial learning phase (d1-3, 4 trial per day, d4, 1 probe trial) mice progressively reduced the number of error (entry in error arm) before finding the target arm (**Fig. 1d left**; *Day effect*; F_(3;33)_=3.202; p=0.0359; see **Supplementary Fig. 1a and 1b** for latency and distance analysis). During the memory probe trial on day 4, mice spent more time in the target arm compared to others despite the absence of a water reward (**Fig. 1d right**; *Arm effect*; F_(4;44)_=7.412; p=0.0001). Thus, mice progressively learned to travel more efficiently toward the location of the target arm. Twenty-five days later, mice continued to show a preference for the target arm position (**Fig. 1e left**; remote memory testing, d28; *Arm effect*; F_(4;32)_=3.766; p=0.0128). On subsequent reinforcement-training days (d29-32) mice did not further increase their performance (**Fig. 1e middle**; *Day effect*; F_(3;24)_=1.529; p=0.2326; see **Supplementary Fig. 1a and 1b** for latency and distance analysis) and displayed a strong preference for the target arm during the subsequent probe test on day 32 (**Fig. 1e right**; *Arm effect*; F_(4;32)_=17.16; p<0.0001). This suggests that mice reached an asymptotic level of performance during this testing phase of reinforcement-training. Finally, mice were tested for reversal learning by switching the target arm to a new allocentric location. During this new learning period (d33-36), mice progressively reduced the distance traveled to reach the new target arm (**Fig. 1f left**; *Day effect*; F_(3;18)_=7.01; p=0.0025; see **Supplementary Fig. 1a and 1b** for latency and distance analysis) and displayed a clear preference for that new location during the day 36 probe trial (**Fig. 1f right**; *Arm effect*; F_(4;24)_=5.417; p=0.0030) suggesting that mice were able to acquire a new spatial memory.

### Temporally correlated MS glutamate neuron activity cluster in two main populations associated with successful navigation toward the target arm

We analysed MS glutamate neuron activity throughout all sessions. Across all testing phases, a total of 103 MS glutamate neurons were detected (**Fig. 2**; n= 12 mice; n=13 recording sessions; and **Supplementary Fig. 2**; Methods). As glutamate neuron activity had previously been associated to locomotion (Fuhrmann et al., 2015; Justus et al., 2017; Zhang et al., 2018), we automatically detected locomotion events (>5cm/s, >1sec; see methods) and examined cell activity around those locomotion events (**Fig.2b**; methods). Across all cells, peak calcium activity spanned heterogeneously around locomotion events (**Fig.2b left**). Using a clustering algorithm to group neurons based on the correlation between their temporal activity patterns around locomotion events, we identified several subpopulations of MS glutamate neurons in accordance to the period where their peak activity occurred (**Fig. 2c**). We used a meta-K-means algorithm (Ozden et al., 2008) employing 5000 runs of K-means++, a seeded version of the traditional K-means algorithm, followed by maximisation of both cluster size and intra-cluster neuron-neuron correlation (Dombeck et al., 2009; Methods). This analysis found two large clusters (cluster one and two; **Fig.2c**) and three smaller clusters that were grouped together (other clusters; **Fig.2c**). The first cluster consists of neurons predominantly active before the onset of locomotion and the second cluster group neurons that had their peak activity near the end of locomotion events (**Fig.2c**). Other clusters did not show any clear activity patterns and displayed a low correlation with both cluster one and two (**Fig.2c**). To assess the cell coding reliability, we ran a split-half correlation analysis (Supplemental Fig.3, methods). Cluster 1 and 2 neurons were more consistent in their activity than cluster 3 neurons (corrected correlation of 0.44 for cluster 1, 0.55 for cluster 2 and 0.14 for cluster 3, Supplemental Fig.3).

Cells from all clusters showed varying correlation with the mice velocity ranging from non-significantly correlated to both positively or negatively correlated (**Supplemental Fig.4a**), suggesting that, in contrast to a previous reports using photometric recordings with slower calcium reporters on head fixed mice running on a treadmill (Fuhrmann et al., 2015; Justus et al., 2017), MS glutamate neuron activity had a more complex relationship with locomotion in freely-moving mice when performing a navigation task. In this navigation task, most of locomotion events began in a given start-arm and ended in a different arm which was either an error arm or the target arm (**Supplemental Fig.5a**). Also, the number of error and target trajectories were comparable for each given training phase (**Supplemental Fig.5d**). MS glutamate neurons were more active in arms than in other part of the maze (**Supplemental Fig.5b-c**). We therefore examined if MS glutamate neuron activity was modulated by the spatial destination of the trajectory.

The activity patterns displayed by neurons in cluster one and two was sorted according to the destination of the trajectory (**Fig. 3**) and whether the destination was correct (leading to the target arm) or incorrect (leading to an error arm). Interestingly, most cells from cluster one (cells more active before locomotion) and two (more active near the end of locomotion) showed significantly higher activity for correct rather than error trajectories. Cells from other clusters did not show this preferential activation (**Fig.3a-c**). The activity pattern of neurons from cluster one was significantly modulated by the spatial destination (error vs. target; n=55 cells Destination * Time interaction; F_(2;216)_=16.99; p<0.0001), with their peak activity occurring preferentially before the onset of trajectories leading to the correct target arm (**Fig.3d top**). Thus, MS glutamate neurons from cluster one that display increased activity before the onset of locomotion appear to code preferentially for future successful trajectory toward the target arm and have been named *preparatory cells*.

**Figure 3.**
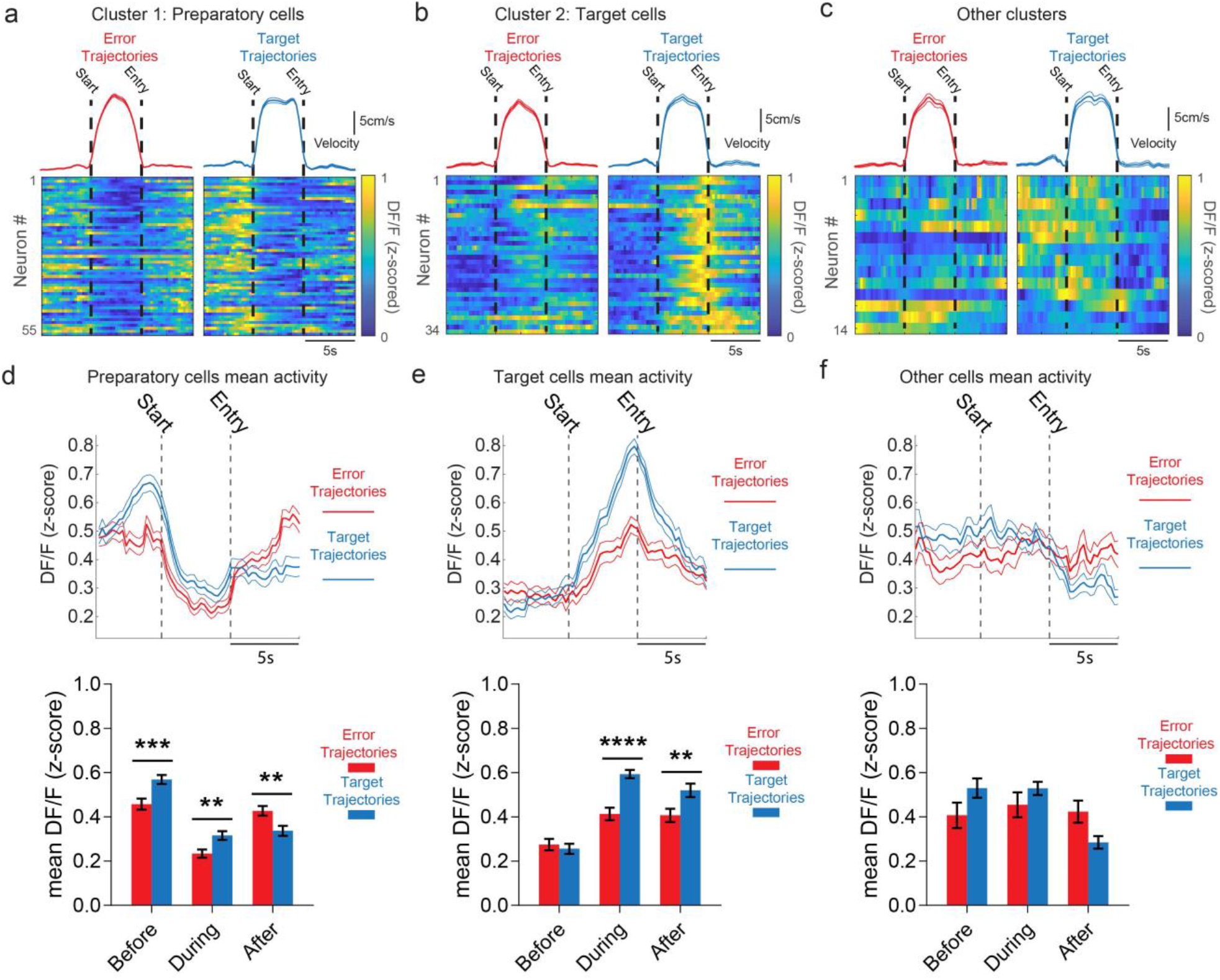
Subpopulation of MS glutamatergic neurons are activated preferentially before locomotion or later in the target zone and are modulated by performance. **a-c**, averaged cell calcium activity and locomotion velocity for trajectories ending into errors and trajectories leading to the target, for the three cell clusters, Preparatory cells (a) Target cells (b) and others (c). Dashed line denotes locomotion onset and arrival into the destination arm. **d-f**, Left panel: averaged calcium trace (± s.e.m.) for error (red) and target (blue) trajectories. Right panel: mean calcium activity per phase (Before locomotion onset, during locomotion, after locomotion), for preparatory cells (d) Target cells (e) and other cells (f). **d**, Preparatory cells were more active before the onset of trajectories leading to the target. **e**, Target cells were more active when mice enter the target arm. **f**, Other cells do not show a preferred phase of activity. Error bars represent s.e.m. *P<0.05; **P<0.01; ***P<0.001; ****P<0.0001 (*Holm-Sidak’s multiple comparison*).

Cluster two neurons were also significantly modulated by the spatial destination (n=34 cells; Destination * Time interaction; F_(2;132)_=9.048; p=0.0002), with an activity that peaked around the entry in the target arm (**Fig.3e**). Thus, cells from cluster two that were preferentially active near the end of the locomotion are specifically activated when locomotion terminates in the target arm but were less active when mice enter an error arm. These cells may therefore code for the entry in the rewarded target arm and were named *target cells*. In contrast, MS glutamate neurons from other clusters were not modulated by the destination (*n=14 Other cells;* Destination * Time interaction; F_(2;52)_=0.1318; p=0.7195) and did not display a clear peak of activity (**Fig.3f**). Mice ran slightly faster during target trajectories than during error trajectories (18.60cm/s vs. 16.54cm/s respectively; t_(102)_=5.8881, p<0.0001). To evaluate the potential influence of locomotion velocity on calcium activity, we used for each mice the median velocity from all locomotion events to sort fast and slow trajectories. This analysis showed that the calcium activity was influenced by the destination but not the locomotion velocity itself (**Supplemental Fig. 4b-d**). Therefore, the slightly faster velocities for target compared to error trajectories may reflect the mice’s motivation to reach the target but alone is unlikely to explain the preferential representation of target trajectories by MS glutamate neurons from cluster one and two. To further examine these two main cell populations, we next investigated how the activity patterns of preparatory and target cells were modulated across the three learning phases.

**Figure 4.**
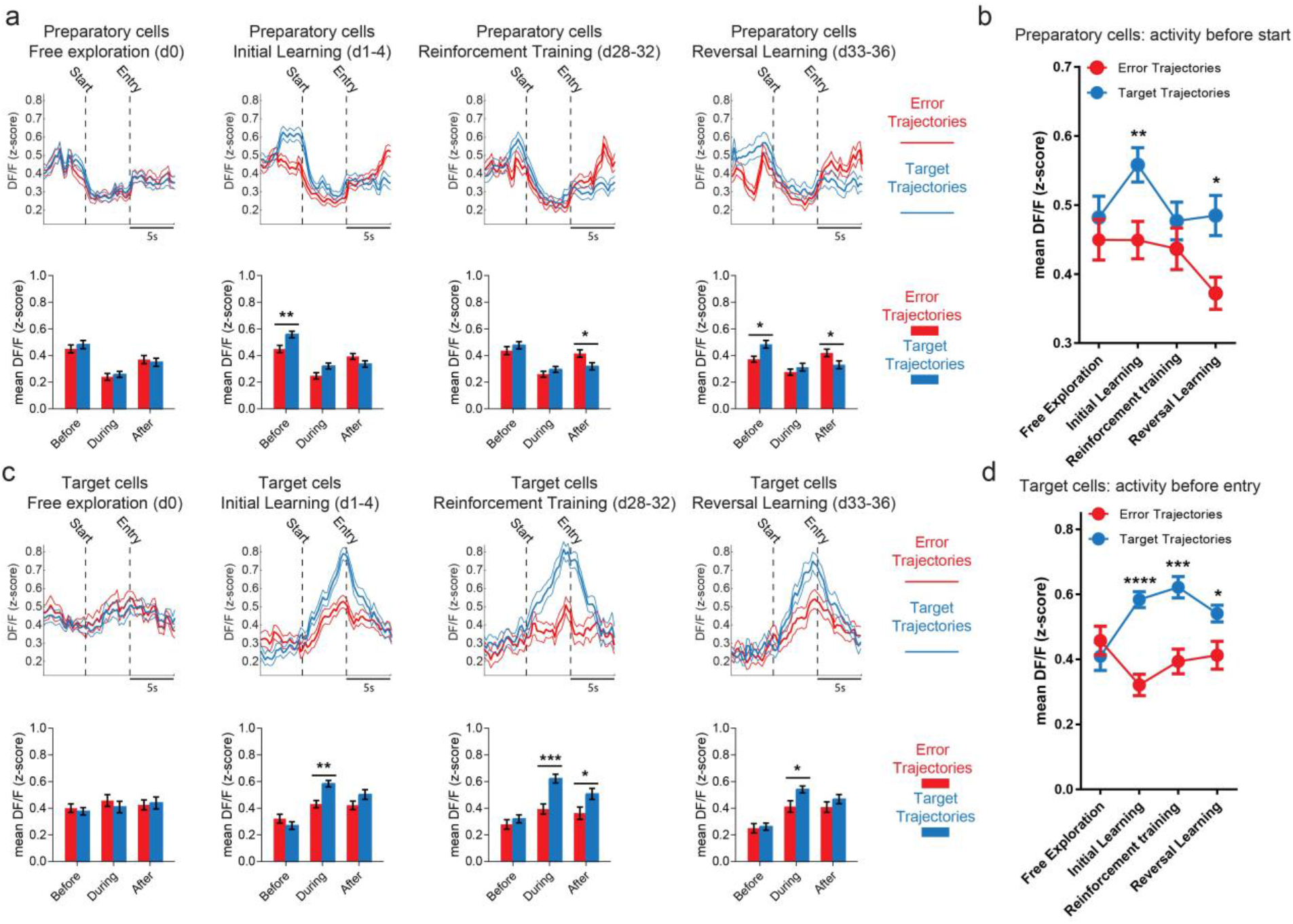
Subpopulation of MS glutamatergic neurons encode differentially target and error trajectories during spatial navigation and learning. **a**, Preparatory cell activity across learning. Far left: Free exploration of day 0. Middle left: Initial learning. Middle Right: reinforcement training. Far right: reversal training. Top: mean cells trace for error and target trajectories (red and blue, respectively). Bottom: mean fluorescence per phase around locomotion. **b**, preparatory cell activity before locomotion onset for all learning phases: preparatory cells are more active for trajectories leading to the target only during the initial and reversal learning. **c**, Target cell activity across learning. Far left: Free exploration of day 0. Middle left: Initial learning. Middle Right: reinforcement training. Far right: reversal training. Top: mean cells trace (± s.e.m.) for error and target trajectories (red and blue, respectively). Bottom: mean fluorescence per phase around locomotion. **d**, Target cell activity during locomotion for all learning phases: target cells are more active for trajectories leading to the target for all phase except the free exploration in with no water reward was provided. Error bars represent s.e.m. *P<0.05; **P<0.01; ***P<0.001 (*Holm-Sidak’s multiple comparison*).

### Learning modulates preparatory and target MS glutamate neuron activity

When naïve mice freely-explored the maze with no reward delivery (Free exploration, d0, Fig.4a), preparatory cells were more active before locomotion (Time effect; F_(2;100)_=35.94; p<0.0001) but did not show increased activity for trajectories toward the future target arm (Fig.4a-b; Destination * Time interaction; F_(2;100)_=0.5734; p=0.5654). In contrast, during the initial learning phase (d1-4, Fig. 4a), *preparatory cells* were more active before the onset of trajectories leading to the target arm compared to error trajectories (Fig.4a-b; Destination * Time interaction; F_(2;176)_=8.836; p=0.0002). During the reinforcement training, preparatory cells continued to be active before locomotion onset (Time effect; F_(2;152)_=27.77; p<0.0001), however, the preferential activation before target trajectory was no longer present (Fig.4a-b; Destination * Time interaction; F_(2;152)_=0.6913; p=0.7933) even though mice reached their asymptotic level of performance during this phase (d28-32; Fig.4a-b). Interestingly, when mice had to learn a new position for the target arm in the last learning phase (d33-36; Reversal learning, Fig.4a), preparatory cells again showed higher activation before target trajectories compared with trajectories leading to errors (Fig.4a-b; Destination * Time interaction; F_(2;132)_=10.36; p<0.0001). As for MS glutamate target cells, they also did not show any specific activity when naïve mice freely-explored the maze (Free exploration, d0; **Fig.4c-d**). However, these cells displayed a stronger peak activation when mice entered the target arm compared to error arms during initial learning (d1-4; Destination * Time interaction; F_(2;104)_=5.953; p=0.0036), reinforcement training (d28-32; Destination * Time interaction; F_(2;60)_=3.958; p=0.0496) and reversal learning (d33-36; Destination * Time interaction; F_(2;76)_=2.009; p=0.0302).

These results show that *MS glutamate preparatory cells* are preferentially activated before target trajectory onset and are strongly modulated by the correct destination in the maze, only when mice were acquiring a new spatial target (initial learning and reversal learning). This activity dependant modulation by the destination was not present when mice had previously learned to navigate to the target position (Reinforcement training). In contrast, *MS glutamate target neurons* showed specific activation when entering the target arms across all learning phases. Therefore, *MS glutamate preparatory cell* activity appear to be required for an efficient navigation toward the target and may facilitate the planning of goal-directed navigation, especially when spatial memory is under formation, while *MS glutamate target cells* present a more stable code for entry in the goal area.

### *MS glutamate preparatory cell* activity is required for spatial goal-directed learning and memory

We hypothesized that the activation of *MS glutamate preparatory neurons* during the initial learning phase was important for spatial learning and memory. To examine if MS glutamate preparatory cell activity is required for spatial learning, we optogenetically silenced MS glutamate neurons at the time when preparatory cells were primarily active. To specifically inhibit MS glutamate neurons, we transfected the MS in VGLUT2-CRE animals using the CRE-dependant AAVdj-flex-ArchT virus (see methods). Virus expression spanned across most of the MS with neurons transfected across both the medial and lateral portions of the MS (**Fig. 5b, Supplemental Fig. 7**). During initial learning, we optogenetically silenced MS glutamate neuron specifically during the start arm waiting portion of the task up until mice entered the first choice point to disrupt their preparatory coding activity (**Fig. 5c, Supplemental Fig. 9**). On day 0, both ArchT and eYFP control mice freely-explored the maze (d0) and displayed no preference for the future target arm (**Fig. 5d**; *Experimental group*destination*; F _(4, 100)_ = 1.017; p = 4024). Across the three day learning period, when optogenetic silencing was restricted during the 30 s start arm period and up to the first choice point, ArchT mice displayed no significant improvement in number of errors to the target across days compared to eYFP controls (**Fig.5e**; *experimental group effect* F_(4, 100)_=16.564; p=0.0168), suggesting that optogenetic inhibition of MS glutamate neurons preparatory activity induced deficits in spatial learning. On day 4, spatial memory was assessed by comparing the amount of time spent in the target arm during the probe trial compared to the other arms. During the probe test, ArchT mice showed no preference for any arm, while eYFP controls had a significant preference for the target arm (**Fig.5e**; *experimental group * arm interaction*; F _(4, 100)_ = 4.319; p = 0.0029), spending more time in the target arm compared to the other arms despite the absence of water delivery. This demonstrates that silencing MS glutamate preparatory activity during the start-arm period results in significant learning deficits, suggesting that *MS glutamate preparatory neurons* play an important role in spatial memory formation. Next, to examine if silencing MS glutamate activity had any influence on free exploration or general locomotion, we tested the effect of inhibiting this population throughout a series of exploration tasks including a linear track, decagon and an open field environment (**Supplemental Fig. 10**). Across the tasks, silencing MS glutamate neurons had no effect on any locomotion-related parameters tested (**Supplemental Fig. 10**). In a rewarded linear track, there was no significant difference between ArchT and eYFP control animals with latency to target or average speed (**Supplemental Fig. 10e-j**). Thus, MS glutamate neurons activity is not required for free locomotion during new environment exploration nor for straight locomotion toward a target in a linear environment.

**Figure 5.**
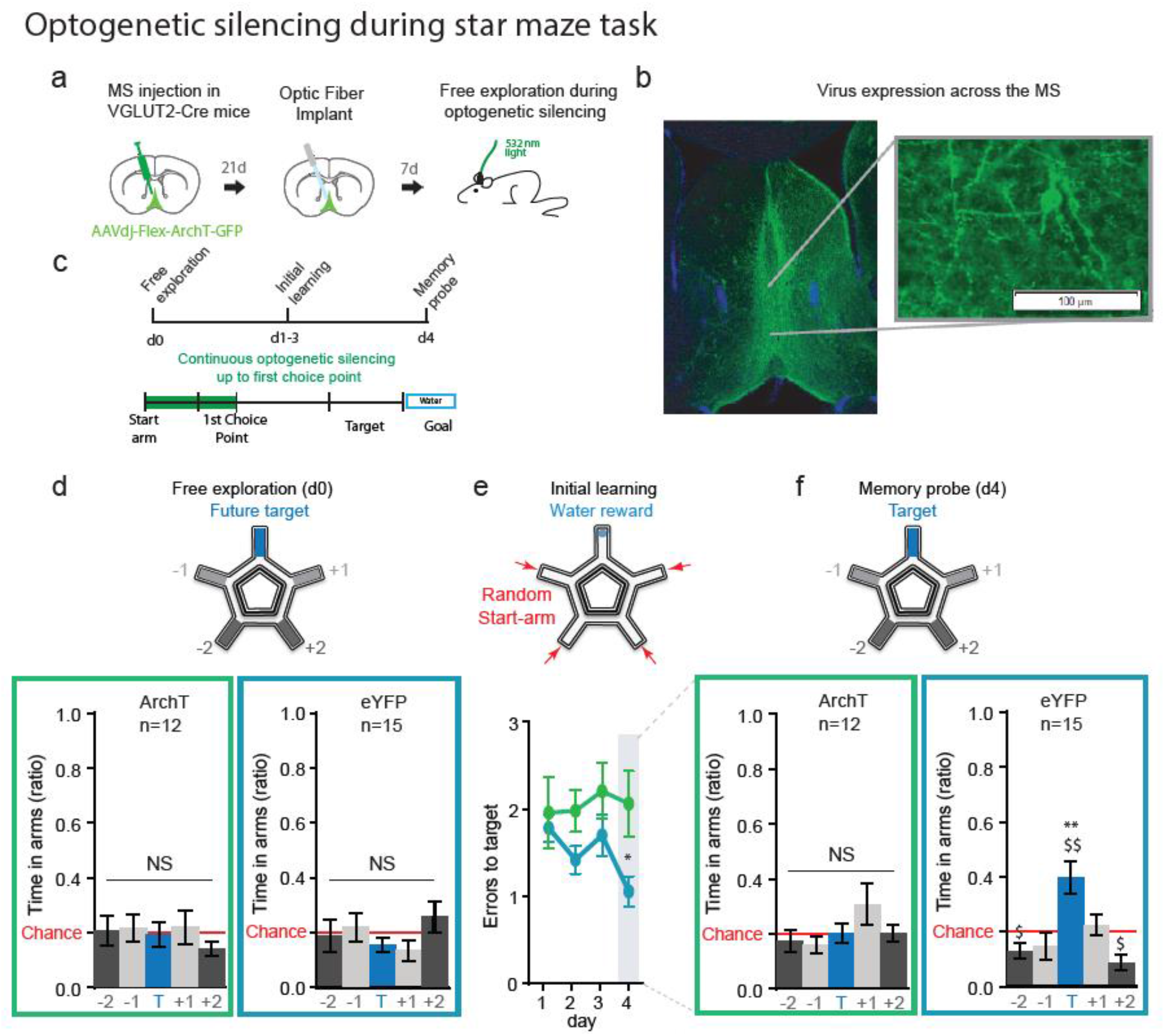
Optogenetic silencing of MS glutamate neurons at the start arm portion of the star maze task reduces memory performance. **a**, Virus and experimental timeline of optogenetic experiments. **b**, AAVdj-flex-ArchT virus expression in the medial septum. **c**, Representation of star maze experiments with optogenetic silencing during 30 second start arm period until first choice point. **d**, Ratio of time spent in each area of the maze, showing no significant preferences for any arm on day 0 (*experimental group*\**Arm interaction*; F _(4, 100)_ = 1.017; P = 0.4024), **e**, Mean errors to target across a 4 day testing period comparing ArchT and control eYFP groups (*experimental group effect* F_(1, 25)_ = 6.564, p=0.0168)). **f**, amount of time spent in the target arm during the probe trial. ArchT animals showed no arm preference compared to controls (*experimental group * arm interaction*; F_(4, 100)_=4.319; p=0.0029). ArchT animals had no preference for the target arm compared to chance (Holm-Sidak’s multiple comparisons; p>0.05) compared to controls (*Holm-Sidak’s multiple comparisons; p=0.0055*). Error bars represent s.e.m. *P<0.05; **P<0.01; (*Holm-Sidak’s multiple comparison*). $P<0.05; $$P<0.01 (One-sample t-test vs. chance).

In summary, taking into consideration the optogenetic and calcium imaging results reported here, in a complex environment where animals must take navigation decision, MS glutamate neurons code for information regarding the spatial destination, and *MS glutamate preparatory cells* appear to be crucial for spatial learning and memory.

## Discussion

The hippocampus contains highly plastic neural circuits that are known to be associated with episodic-like memory in humans, a memory that rely on *what-where-when* representations that can be flexibly used. In rodents, episodic-like spatial navigation, learning and memory that are acquired over a few numbers of trials also depend on hippocampal circuits (Dickerson et al., 2010). Depending on the memory processes engaged and on the age of the memory trace, different hippocampal related circuit are recruited (Lux et al., 2016). As the MS is one of the main input pathways to the hippocampus, it is well poised to contribute to these functions. The main purpose of the current study was to investigate the activity of MS glutamate neurons in the freely behaving mouse during an episodic-like spatial learning and memory task, and test how this population contributes to spatial memory.

It has been over a decade since MS glutamate neurons were first identified (Sotty et al., 2003; Sotty et al., 2003) and during this time several lines of research have elucidated their connectivity and possible contributions to the septo-hippocampal network. MS glutamate neurons have been shown to project to both GABAergic and cholinergic neurons within the MS, and project to both interneurons and pyramidal cells in the hippocampus (Manseau et al., 2005, Huh et al., 2010; Fuhrmann et al., 2015; Robinson et al., 2016). MS glutamate neurons also project across multiple layers of the entorhinal cortex (Manns et al., 2001; Gonzalez-Sulser et al., 2014; Justus et al., 2017). Previously, it has been shown that rhythmic activation of MS glutamate neurons results in the driving of hippocampal rhythms across theta frequencies, indicating that they may participate in locomotion and rhythm generation (Fuhrmann et al., 2015; Robinson et al., 2016). The data presented here suggests that their role is more complex, with MS glutamate neurons forming a heterogeneous population that code for multiple aspects of the environment including spatial navigation planning and target or reward location.

Recent evidence indicates that MS glutamate neurons are related to locomotion initiation since optogenetic activation of these neurons increase movement (Fuhrmann et al., 2015; Zhang et al., 2018) and the activity of MS glutamate neuron terminals in the entorhinal cortex are modulated by speed (Justus et al., 2017). In this study, we characterized the spontaneous activity of MS glutamatergic neurons using miniature endoscopes to perform calcium imaging with single-cell resolution during a star-maze episodic-like spatial learning task to identify activity-associated changes during learning. We then examined the effect of optogenetic manipulation and found causal evidence for the contribution of MS glutamate cells to navigation and spatial memory.

We found a heterogeneity of MS glutamate neuron activity tightly associated with different sections of the star maze and learning states. Two main MS glutamate functional subpopulations were identified. The first population of glutamate neurons were primarily active in the start-arm before locomotion onset, identified here as *MS glutamate preparatory neurons*. These neurons had significantly higher activity before trajectories leading to the target arm compared with trajectories leading to error arms. This increase activation was only present when there was a learning component to the task but did not show any arm specific activation when the animal was freely exploring. Interestingly, this activity was positively-modulated before mice engaged in locomotion leading to the target only when the animal had to acquire a new memory (initial learning and reversal learning phases) but not when mice navigated in a well-known goal (retraining phase). This suggests that the activity of preparatory MS glutamate neurons could contribute to selecting, planning and/or retrieving the appropriate route to the rewarded target, especially when memory is still labile (new learning) but not when memory is highly consolidated (remote memory during retraining). It is possible that activation of MS glutamate preparatory neurons before navigation initiation may facilitate the retrieval of hippocampal place cell assemblies coding for the correct trajectory toward the target arm (Ainge et al., 2007), allowing for the mice to select the appropriate path to this goal location. With regards to such retrieval of the correct hippocampal representation, it is interesting to note that the terminals of MS glutamate neurons are known to activate OLM interneurons in the hippocampus (Fuhrman et al., 2015; Robinson et al., 2016), an interneuron type proposed to enhance CA3 input strength within CA1 through local circuit modulation (Leao et al., 2012). Such a MS glutamate-OLM circuit, through their enhancement of CA3 input, could therefore favour recall which is important for planning navigation strategies before initiating the trajectory. In this context, the absence of target-specific preparatory activity in MS glutamate neurons when mice reached their optimal level of performance (remote memory during the retraining phase) may be explained by the fact that CA3-dependant memory may be less required for retrieval of a remote memory compared to a more recent labile memory trace (Lux et al., 2016). Alternatively, MS preparatory neurons may also be important for goal-driven attentional processing such as is observed in neurons of the cortex (Kim et al., 2016). This phenomenon of preparatory neurons activity has been observed previously in the pre-motor and motor cortices (Economo et al., 2018), in visual and prefrontal cortices, and in subcortical regions such as the striatum (Schultz et al., 2003) and substancia nigra dopamine neurons (Da Silva et al., 2018). However, our results suggest that *MS glutamate preparatory neurons* may not be directly related to locomotion *per se* as silencing of MS glutamate cells in the start-arm of a linear track did not affect locomotion speed and the latency to target ruling out a direct effect on movement initiation. In contrast, *MS glutamate preparatory neurons* silencing during the start-arm waiting portion of the star maze task (before locomotion initiation), induced a dramatic impairment in spatial memory learning and retrieval. Taken together, these results indicate that *MS glutamate preparatory neurons* are key contributors to efficient goal-directed navigation planning when recent spatial memory is still labile.

The second population, *MS glutamate target neurons* had their activity peak around the end of the locomotion when mice entered the target arm. Interestingly, the activity of these neurons was strong during all learning phases (initial learning, reinforcement and reversal learning) which suggest they likely play a role in signalling reward proximity. Activation of those target MS glutamate neurons was dependant on the presence of a reward as they did not show any activation during the free-exploration session of the maze. Thus, these cells appear to code for the proximity of the reward-associated spatial context. Such reward-proximity signal may be important for driving goal-directed responses as MS glutamate neurons are known to project to the ventral tegmental area (Fuhrmann et al., 2015). Unfortunately, that subpopulation of MS glutamate neurons was not optogenetically targeted as optogenetic inhibition starting at the vicinity of the target arm may represent an external cue helping the mice to detect the target arm. The activity of these neurons bears some resemblance to neurons signalling reward-related information in the midbrain dopamine system that are activated in the presence of a reward-predicting stimulus (Schultz 2007). It has been recently shown that MS activation increases the activity of dopaminergic neurons from the ventral tegmental area while inhibiting the activity of dopaminergic neurons from the substancia nigra (Bortz and Grace 2018). Such MS modulation has been proposed to attenuate substancia nigra-driven impulsive response favouring VTA-driven goal-directed response (Bortz and Grace 2018). Although these effects have been associated with cholinergic and GABAergic septo-hippocampal projections, it is unclear how MS glutamate neurons modulate dopaminergic neuron activity. Such modulation may be elicited through direct MS glutamate projection to the VTA (Fuhrmann et al., 2015), through septo-hippocampal glutamate projection or following intra-MS glutamate collaterals to both GABA and cholinergic neurons (Manseau et al., 2005, Huh et al., 2010; Fuhrmann et al., 2015; Robinson et al., 2016). The origin of such target signal in MS glutamate neurons remain unclear but may arise from both the VTA itself (Aransa et al., 2015) or from lateral hypothalamic area, a region implicated in feeding and reward (Stamatakis et al., 2016) that project directly to MS glutamate neurons (Fuhrmann et al., 2015). Disentangling the role of these networks will require further investigations.

Taken together, in addition to coding for the locomotion velocity itself, MS glutamate neurons encode cognitive information about the spatial destination in the environment. MS glutamate neurons play an essential role throughout the learning phase of the task and silencing these neurons during the preparation phase of navigation induces significant deficits in task performance. This points towards MS glutamate neurons playing an essential role in trajectory planning, learning or memory encoding. Future experiments will aim to explore their exact contribution to learning and retrieval. These findings represent a major gain in the understanding of spatial learning and memory with important potential mental-health related implications as the MS is dramatically altered early in the pathological course of memory-related disorders such as Alzheimer Disease.

## Acknowledgment

this work was supported by the Fyssen foundation (JB.B.) CIHR, NSERC and Brain Canada

## METHODS

### Virus injections

All procedures were performed according to protocols and guidelines approved by the McGill University Animal Care Committee and the Canadian Council on Animal Care. VGLUT2-Cre knock-in homozygote mice (The Jackson Laboratory, stock #016963) were housed in a 12:12 hour light/dark cycle with food and water ad libitum. To trigger the expression of the opsins, VGLUT2-Cre mice were stereotaxically injected in the septum with a Cre-dependent adeno-associated viral vectors. Activation experiments were performed with ChETA-eYFP (AAVdj-Ef1-DIOhChR2(E123T/T159C)-eYFP from Molecular Virology Support Core, Oregon Health and Science University or Stanford Neuroscience Gene Vector and Virus Core, Stanford University). Silencing experiments were performed with ArchT-GFP (AAV2-EF1α-Flex-ArchT-GFP from University of North Carolina Virus Core). Calcium imaging experiments were performed with a Cre-dependant GCAMP6f reporter (AAV9.Syn.Flex.GCaMP6f.SV40, from Penn Vector Core, University of Pennsylvania). Control experiments were performed with Cre-dependent eYFP control virus (AAVdj-Ef1-Flex-eYFP from Vollum Vector Core, Oregon Health and Science University). For all optogenetic experiments, adult mice were injected at approximately 10 weeks of age, and for all calcium imaging experiments, mice were injected at approximately 18 weeks of age. Mice were anesthetized using isoflurane and positioned in a stereotaxic frame (Stoelting), and viruses were delivered into the septum (0.6 μl at 0.06 μl/min). Animals were injected as follows: AP 0.86 mm from bregma, ML 0.5 mm, DV 4.5 mm at a 5° angle.

### Calcium Imaging

Mice were injected at approximately 18 weeks of age and following a 3-4 week incubation period, were implanted with 8 by 0.5 mm diameter GRIN lens to be imaged with a head mounted miniaturized fluorescence microscopes system (Doric Lens, Quebec, Canada). GRIN lens placement was implanted at a 5° angle into the medial septum (AP 0.86, ML 0.2). Lens depth was set based on fluorescent neurons visible in the field of view during the implantation, and ranged between 3.8 mm and 5 mm deep. The baseplate was secured to the skull using metabond and dental cement (Patterson Dental). Following a 3-week recovery period, animals were water restricted to 1.5 hrs of water availability per day to improve the quality of the recorded individual cell resolution. During recordings, GCaMP6f signal was visualized with 0.6 mW power (4,5mW/mm2 intensity) of 458 nm light and videos were sampled at 20 fps. Behavioural events where synchronized with imaging by acquiring a TTL output from the Miniscope acquisition box to the behavioural recording system (Neuralynx).

### Optogenetic Experiments

Mice were injected at ~10 weeks of age and, following a 3-4 week incubation period, were implanted with an optic fiber (Thorlabs) connected to a ferrule (Precision Fiber Products), which was implanted at a 5° angle to target just above the medial septum (AP 0.86, ML 0.2, DV 3.83 mm). Headstage and optic implants were secured to the skull with metabond and dental cement (Patterson Dental). Following surgery, animals had 1-week of recovery followed by a 1-week habituation period. Once habituated, animals were connected to an optic fiber patch cord (Thorlabs). Light delivery was achieved through an optic fiber patch cord and sleeve coupled to a 530 nm laser (LaserGlow), and light intensity at the tip of the optic fiber was set for 20 mW.

### Behavioural testing in a modified allocentric Star-maze

The Dry Star-Maze task (DSM) test was developed in the laboratory to assess spatial reference memory in mice over 5 consecutive days (adapted from Rondi-Reig et al., 2006). Both arms and alley were 20 cm long. In the DSM, water-restricted mice freely explore the device until they find the only baited arm (target arm) containing 0.15ml of water mixed with 9% sucrose. Each day began with a baseline recording of 60 seconds (home-cage). At the end of the baseline, the mouse is confined in a randomly–selected start-arm for 30 seconds and is then free to explore the maze (cut-off: 5 minutes). Between trials, mice were placed back in a familiar box for 1 minute ITI. During ITI, the maze was cleaned with 70% ethanol to disrupt odour-cues based strategy. The maze was wiped clean and pseudo-randomly rotated between mice to favour allocentric strategies. On day 0, mice were habituated to the environment during a freely exploration trial for 5 minutes. During free exploration, if a spontaneously preferred arm was identified, this arm was not selected as the target arm in subsequent learning phase. Calcium imaging experiments were divided into three learning phases. During the initial learning phase, mice had to learn how to localise the target arm, 25 days after a retraining occurs to examine the contribution of MS glutamate neuron to remote spatial memory and finally, during the reversal learning phase mice had to learn a new target arm position in order to examine contribution of MS glutamate neurons to spatial memory flexible updating. Each learning phase consist in 4 days of testing. The target was reinforced for each mouse from days 1 to day 3 with each day consisting of 4 trial per day (5 minutes cut-off; 60s ITI). At the beginning of each trial, the mouse was confined for 30 seconds in the start-arm and then was allowed to freely explore the maze. If the mouse found the target within the 5 minutes, the trial end, the mouse is allowed to drink the 0.15 ml of sucrose and then is placed back in the neutral cage (beginning of the ITI). If the mouse did not find the target after 5 minutes, the mouse was gently guided to the target. Each trial start from a pseudo-randomly selected start-arm, all non-target arm being used one time each day. For optogenetic experiments, glutamatergic silencing was restricted to the 30 second wait period in the target arm until the mouse reached the first choice point. During the learning phase, the number of errors (entries into a non-target arm), the distance run, and the latency to find the target were measured to assess spatial learning. Memory performance was evaluated on day 4 during one probe-trial without water-reinforcement. In addition, during the probe test, the total number of entries in each arm, as well as the time spent in each arm, were measured to evaluate memory performance.

### Behavioural testing for optogenetic experiments

Optogenetic experiments included four randomized behavioural paradigms including two free exploration tasks and two goal directed tasks. Free exploration tasks include: (1) open field exploration (44 x 44 cm square box with a 30 cm high walls), (2) decagon open field (diameter 93 cm, no walls). Goal directed tasks include: (3) linear track task (LTT) and (4) spatial reference memory star maze task. Open field and decagon exploration tasks consisted of 2-minute free exploration in the novel environments. With optogenetic experiments, glutamatergic neurons were silenced for the entire 2-minute period. The LTT was first used to assess free exploration with 2 minutes in the novel environment. The total distance as well as the latency to reach end of the arm was measured. For each trial, the mouse was confined for 30 seconds in the start-arm before freely exploring the environment. Following free exploration, animals were water restricted and navigate towards 0.15 ml of water mixed with 9% sucrose placed at the end of the linear track. Three trials were performed and between trials, mice are placed back in their familiar box for 1-min (Inter-Trial-Interval; ITI). The water restricted LTT was repeated for two to three days consecutively to habituate animals to receiving water. For LTT optogenetic experiments, glutamatergic neurons were silenced for the 30 second start arm period.

### Anatomical characterization

To examine the expression of the opsins, immunocytochemistry was used to assess virus distribution across the septum. For these experiments, virus injected animals were euthanized at 4– 5 weeks following injections. Mice were perfused and sectioned as described previously (Amilhon et al., 2015). Free-floating sections across the septum were incubated in GFP rabbit antiserum (1:1000, Invitrogen), and were detected with anti-rabbit coupled to AlexaFluor-488 (1:1000, Invitrogen). The slices were mounted with Fluoromount-G (Southern Biotechnology) and analysed with an AxioObserver.Z1 microscope (Carl Zeiss).

### Calcium imaging analysis

Calcium movies were spatially down sampled by a factor of 4 and resulting stacks were motion-corrected with ImageJ (NIH, USA) using the plugging MOCO (https://github.com/NTCColumbia/moco). Then, cell segmentation and calcium trace extraction was performed using the PCA-ICA algorithm as previously described (Mukamel et al., 2009) using custom Matlab scripts (MathWorks). The raw signal is first converted as DF/F to extract fluorescence fluctuation. Then Principal Component Analysis (PCA) was computed for each pixel variance followed by an Independent Component Analysis (ICA), which allowed for the cell extraction. Due to the very low cell density (never more than 11 cells per field of view per day) in line with MS glutamate neurons density, cells were registered across day using the minimal footprint centroid distance followed by a manual verification for all movies. To correct for out-of-focus contamination, for each region of interest (ROI) we subtracted the average fluorescence from a 20μm ring around that ROI. Decreases in baseline fluorescence due to GCaMP6f bleaching were corrected using a 300s-sliding window mean filtering. Average fluorescence trace for each cell was expressed in ΔF/F where F is the average fluorescence for that given cell across the whole recording session. For comparison between cells and across sessions, ΔF/F was then z-scored. For analysis on calcium transients (Supplemental Fig.4), transients were detected as events during which the DF/F trace goes over a 2 standard deviation threshold, and the transient onset frame was determined as the frame the threshold is crossed.

Locomotion events were automatically detected using custom scrip in Matlab. Locomotion was defined as events with a peak velocity that reached at least 5cm/s, locomotion onset and offset were defined using a 2.5cm/s threshold. Locomotion events had to last for at least 1s and two locomotion events separated by less than 2s were pooled together. To analyse calcium activity regarding to locomotion events, the cell activity trace during each locomotion events were linearly realigned to fit the median round duration of locomotion events across mice (5s). Locomotion events that ended in arms were classified as error or target trajectories based on the type of arm in which locomotion ended.

Cell clustering was done as described earlier (Dombeck et al., 2009) using all cells averaged trace around locomotion events. We employed 5000 runs of a regular K-means that was seeded each run by using the K-means++ seeding strategy. Each run generated 4 clusters. Neurons that were in the same cluster >80% of the runs were grouped in meta-cluster. Final cluster were generated in combining meta-clusters for which the correlation between all neurons from each metacluster was greater that the threshold Tcorr. Tcorr was determined by running several time the meta-K-means clustering with increasing values of Tcorr (0.45-0.95, steps of 0.025). The selected value of Tcorr was the value that maximized both the mean cluster size and the mean of the intracluster neuron-neuron correlation. Internal reliability of each cluster’s cell activity was examined by computing split-half reliability correlation for each neuron by calculating the correlation between averaged odd- and even-locomotion events, corrected using the Spearman-Brown prophecy formula (Nunnally et al., 1967).

#### Patch-clamp recordings

VGLUT2-Cre mice were injected with AAVdj-ArchT or AAV2/9-Flex-GCAMP6f were euthanized at 18–30 days after viral injection. Animals were anesthetized animal with ketamine/xylazine/acepromazine mixture and perfused intracardially with cold NMDG solution (at 4°C). NMDG recovery solution contains the following solution (mM): 93 NMDG, 93 HCL, 2.5 KCl, 1.2 NaH2PO2, 30 NaHCO3, 20 HEPES, 25 Glucose, 5 sodium ascorbate, 2 Thiourea, 3 sodium pyruvate, 10 MgSO4, 0.5 CaCl2. Following brain extraction, the brain was incubated for 1 minute in cold NMDG solution and then sliced on the vibrotome with 350 μM sections. Sections were placed in NMDG solution for 10-15 minutes at 32°C and then transferred for a 1 hr recovery in room temperature aCSF solution. Sections were continuously oxygenated with 95% O2/5% CO2. Coronal sections of the septum (350 μm thick) were sliced and recorded using the same protocol as previously published (Huh et al., 2010). Patch glass micropipettes (Warner Instruments) with a resistance 2.2–5 μM were used, and intrapipette solution contained the following (in mM): 124 NaCl, 2.5 KCl, 1.2 NaH2PO4, 24 NaHCO3, 5 HEPES, 2 MgCl2, 2 Na2ATP, 0.3 GTP, 12.5 Glucose, adjusted to pH 7.4 with KOH. In the recording bath, sections were perfused with aCSF and continuously oxygenated with 95% O2/5% CO2 at 5 ml/min at 25°C. To examine the direct responses of VGLUT2 positive neurons transfected with AAVdj-ArchT-GFP in septal sections, GFP-positive neurons were targeted. Recordings were performed in current clamp while cells were held at −70 mV or just above action potential threshold at approximately −55 mV. Silencing was achieved using orange light (532 nm) with a custom-made LED system, consisting of LEDs (Luxeon) coupled with 1-mm-diameter light guide for light delivery (Edmund Optics). Cells were first characterized in current clamp, with current steps of 20 pA to depolarize and hyperpolarize the cell to characterize firing frequency. Following this, 500 ms pulses of light were applied in both current clamp and voltage clamp to assess the hyperpolarization and photocurrent elicited by light. To examine the response of light in an active cell, 1 ms and 30 second square pulses of light were applied repeatedly with cells held above threshold. To characterize calcium imaging responses neurons transfected with AAV2/9-Flex-GCAMP6f cells were recorded in cell-attached mode and spontaneous activity was captured with electrophysiological recording combined with video of calcium responses.

### Patch-clamp data analysis

Patch-clamp recordings were analysed using Clampfit 9 (Molecular Devices). Electrophysiological recordings *in vivo* were analyzed using custom MATLAB scripts (The MathWorks). Spectral analysis was performed using a multitaper fast Fourier transform (Chronux package, window 5 s, step 0.5 s, tapers [1, 1]) (Bokil et al., 2010). Theta peak frequency and peak power (power at the peak frequency) were extracted from the power spectrum. Oscillation strength was used as an index of rhythmicity and was derived from the autocorrelogram (peak value of the power spectrum of the autocorrelation). For all recordings, 2 to 5 repetitions of 10 seconds segments were extracted for analysis. Data was filtered off-line at 3–55 Hz, and spectral analysis was performed using window size 2 seconds, step size 0.5 s. Changes in power, frequency and rhythmicity were compared with baseline values.

### Statistical analysis

Statistical analysis was performed using Prism 6 (Graph Pad). In all figures, data are mean ± SEM. calculated across mice except for calcium analysis, in which the sampling unit was the cell. Multi-factorial ANOVA (factor optogenetic-group, time) and ANOVA with repeated measures (days and arms for the behaviour) were performed. In case of significant interaction among factors, multiple comparisons among groups were performed using Holms-Sidak post hoc test correcting for multiple comparison. Comparison to chance level in the Star-maze was done using a one-sample Student’s t-test. The threshold for statistical significance was set at p=0.05. The experimenter was blinded to the experimental group identity during optogenetic experiments.

## Notes

### Competing Interest Statement

The authors have declared no competing interest.

