## Supplemental material for "Medial septum glutamate neurons are essential for spatial goal-directed memory"

### SUPPLEMENTAL FIGURE AND LEGENDS.

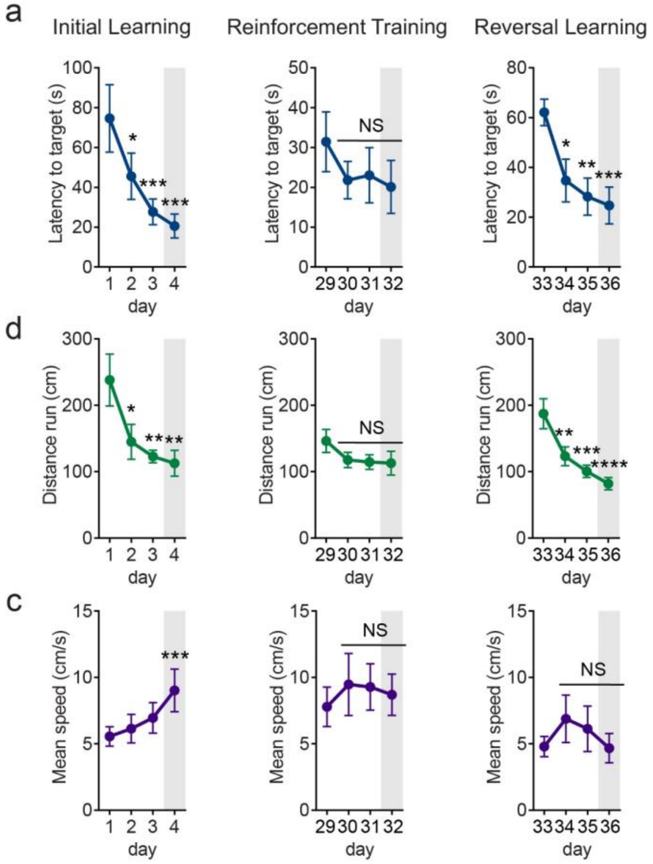


**Supplemental Figure 1 | Freely-moving $Ca^{2+}$ imaging of medial septum glutamatergic neurons during a spatial navigation learning task. a**, Latency to target. Left, initial learning: across learning days, mice reduced the latency before entering the target arm (*Day effect; F(3,33) = 8.8991; p=0.0002*). Middle, reinforcement training: across retraining, mice did not further reduce the latency before finding the target arm (*Day effect; F(3,24) = 0.9725; p=0.4220*). Right, reversal learning: during reversal learning mice progressively reduced the time needed to fin the target arm (*Day effect; F(3,18) = 8.2000; p=0.0012*). **b**, Distance run to target. Left, initial learning: across learning days, mice reduced the distance run before entering the target arm (*Day effect; F(3,33) = 5.153; p=0.0049*). Middle, reinforcement training: across retraining, mice did not further reduce the distance run before finding the target arm (*Day effect; F(3,24) = 0.1847; p=0.1847*). Right, reversal learning: during reversal learning mice progressively reduced the distance run before entering the target arm (*Day effect; F(3,18) = 14.46; p<0.0001*). **c**, Mean trial velocity. Left, initial learning: across learning days, mice progressively increased their mean velocity (*Day effect; F(3,33) = 7.663; p=0.0005*). Middle, reinforcement training: across retraining, mice did not further increase their mean velocity (*Day effect; F(3,24) = 0.46; p=0.7128*). Right, reversal learning: mice did not significantly change their mean velocity during reversal learning (*Day effect; F(3,18) = 1.419; p=0.2700*). Error bars represent s.e.m. *P<0.05; **P<0.01; ***P<0.001; ****P<0.0001 (*Holm-Sidak's multiple comparison*).

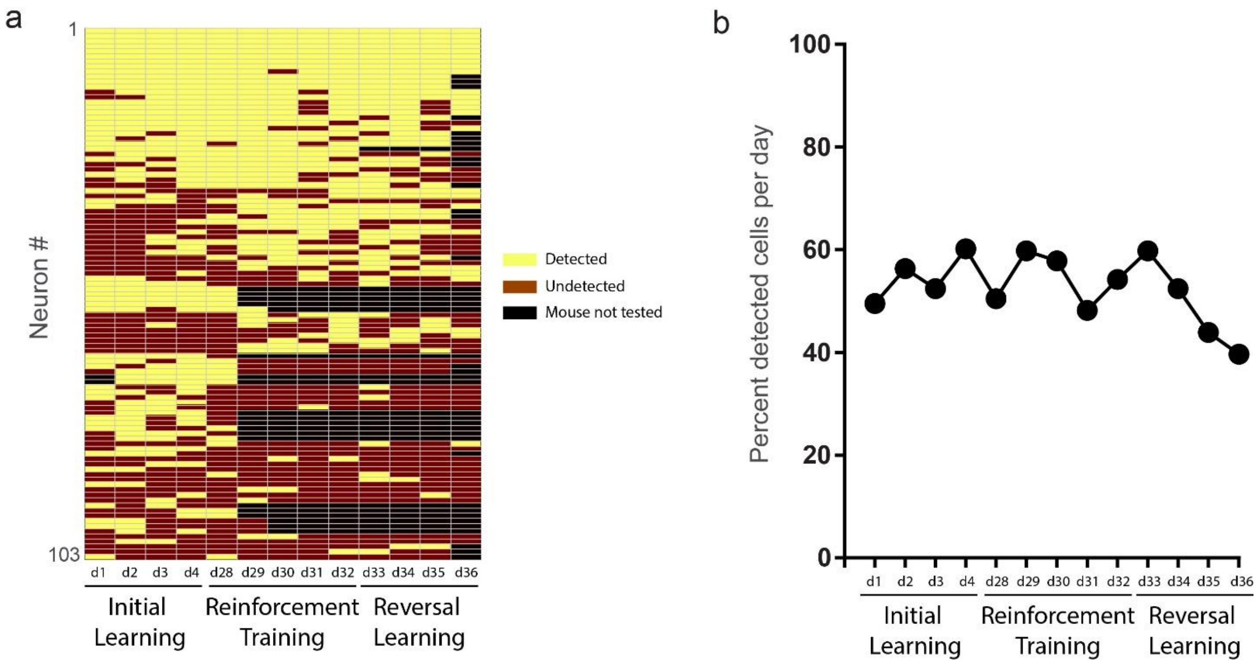


**Supplemental Figure 2 | Cell presence across days. a**, Total number of cells detected per testing day. Each row represents a MS glutamate neuron, and each colon represents a testing day. MS glutamate neurons were not all detected throughout all testing days. Yellow, cell detected; dark red, cell not detected; black, mice not tested. **b**, Fraction of possible cell detected per testing day. Across all sessions, around 50% of all cells were detected each day.

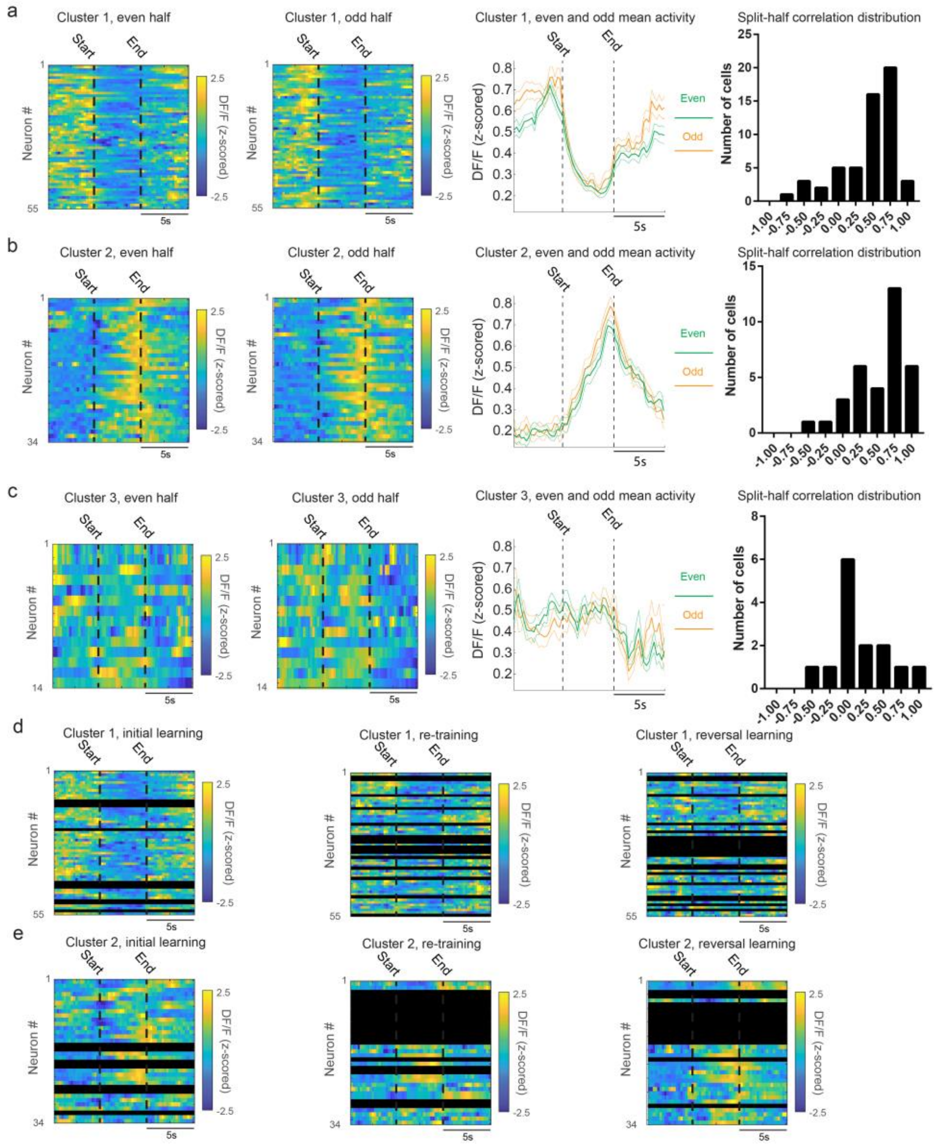


**Supplemental Figure 3 | Split-Half correlation analysis. a**, Cluster 1 split-half correlation analysis. Left, realigned mean activity around even and odd locomotion events for cluster 1 neurons. Right: mean trace for cluster 1 neurons for both even and odd locomotion events. Far right: histogram distribution of cluster 1 neuron internal consistency correlation values. **b**, Cluster 2 split-half correlation analysis. Left, realigned mean activity around even and odd locomotion events for cluster 2 neurons. Right: mean trace for cluster 2 neurons for both even and odd locomotion events. Far right: histogram distribution of cluster 2 neuron internal consistency correlation values. **c**, Cluster 3 split-half correlation analysis. Left, realigned mean activity around even and odd locomotion events for cluster 3 neurons. Right: mean trace for cluster 3 neurons for both even and odd locomotion events. Far right: histogram distribution of cluster 3 neuron internal consistency correlation values. **d**, Cluster 1 active cells average activity for each training phase: initial learning (left), re-training (middle) and reversal-learning (right). **e**, Cluster 2 active cells average activity for each training phase: initial learning (left), re-training (middle) and reversal-learning (right). Error bars represent s.e.m

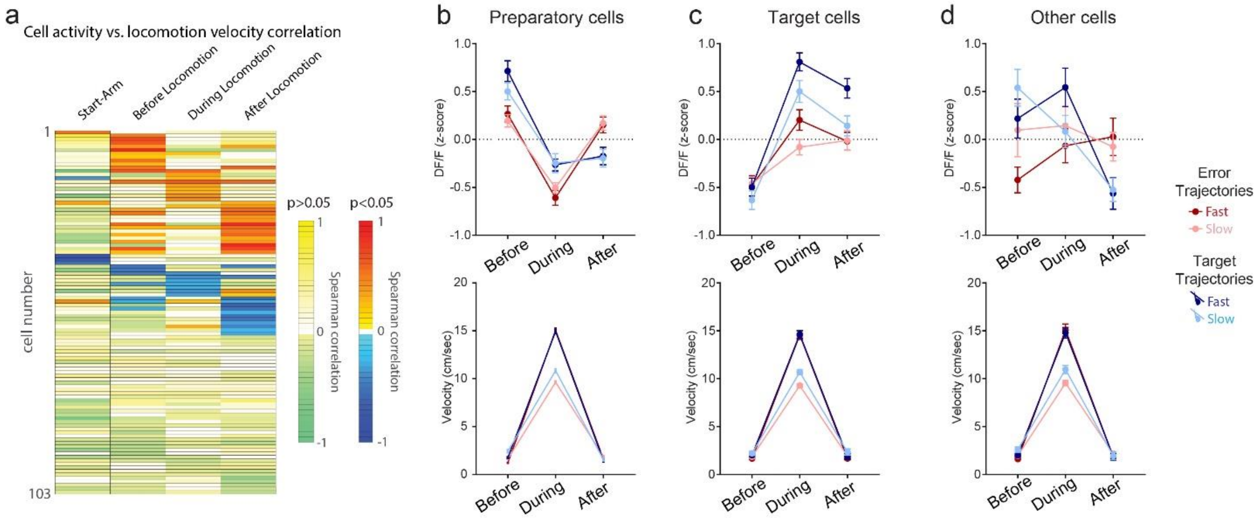


**Supplemental Figure 4 | Velocity-related analysis. a**, Inconstant correlation between cell activity and velocity during locomotion. Overall, 45 out of 103 MS glutamate neuron (43.69%) did not display any significant correlation with locomotion velocity, 35 MS glutamate neuron (33.98%) displayed a positive correlation with locomotion velocity and 23 (22.33%) were negatively correlated with mice velocity. Each row represents the spearman correlation coefficient between a single neuron DF/F trace amplitude and the mean velocity during locomotion, with colors representing the spearman correlation for both no significative correlation (left scale bare) and significant correlation (right scale bare). Each colon represents the mean calcium activity during the start-arm waiting period, during 5s before locomotion onset, during the locomotion and during 5s after locomotion offset. MS glutamate neuron are sorted based on the strength of the correlation with velocity, then by the direction of that correlation (positive correlation, then negative correlation). **b, c and d**, Fast vs. velocity analysis. To determine the influence of locomotion velocity on MS glutamate neuron activity, locomotion events were split based on the medial locomotion velocity. Then, locomotion events were sorted based on the destination (error arm vs target arm). **b**, Preparatory cells are modulated by destination, not by velocity. Top: Preparatory cells activity was significantly influenced by the destination (*Destination * Time interaction* $F(2,2) = 23.11; p<0.0001$) but not the velocity (*Velocity * Time interaction*; $F(2,2) = 1.593; p=0.2041$). Bottom: mean velocity for all trajectory types. **c**, Target cells are modulated by destination, not by velocity. Top: Preparatory cells activity was significantly influenced by the destination (*Destination * Time interaction* $F(2,2) = 12.65; p<0.0001$) but not the velocity (*Velocity * Time interaction*; $F(2,2) = 1.489; p=0.2268$). Bottom: mean velocity for all trajectory types. **d**, Other cells are modulated by destination, not by velocity. Top: Preparatory cells activity was significantly influenced by the destination (*Destination * Time interaction* $F(2,2) = 8.812; p=0.0002$) but not the velocity (*Velocity * Time interaction*; $F(2,2) = 2.46; p=0.0888$). Bottom: mean velocity for all trajectory types. Error bars represent s.e.m.

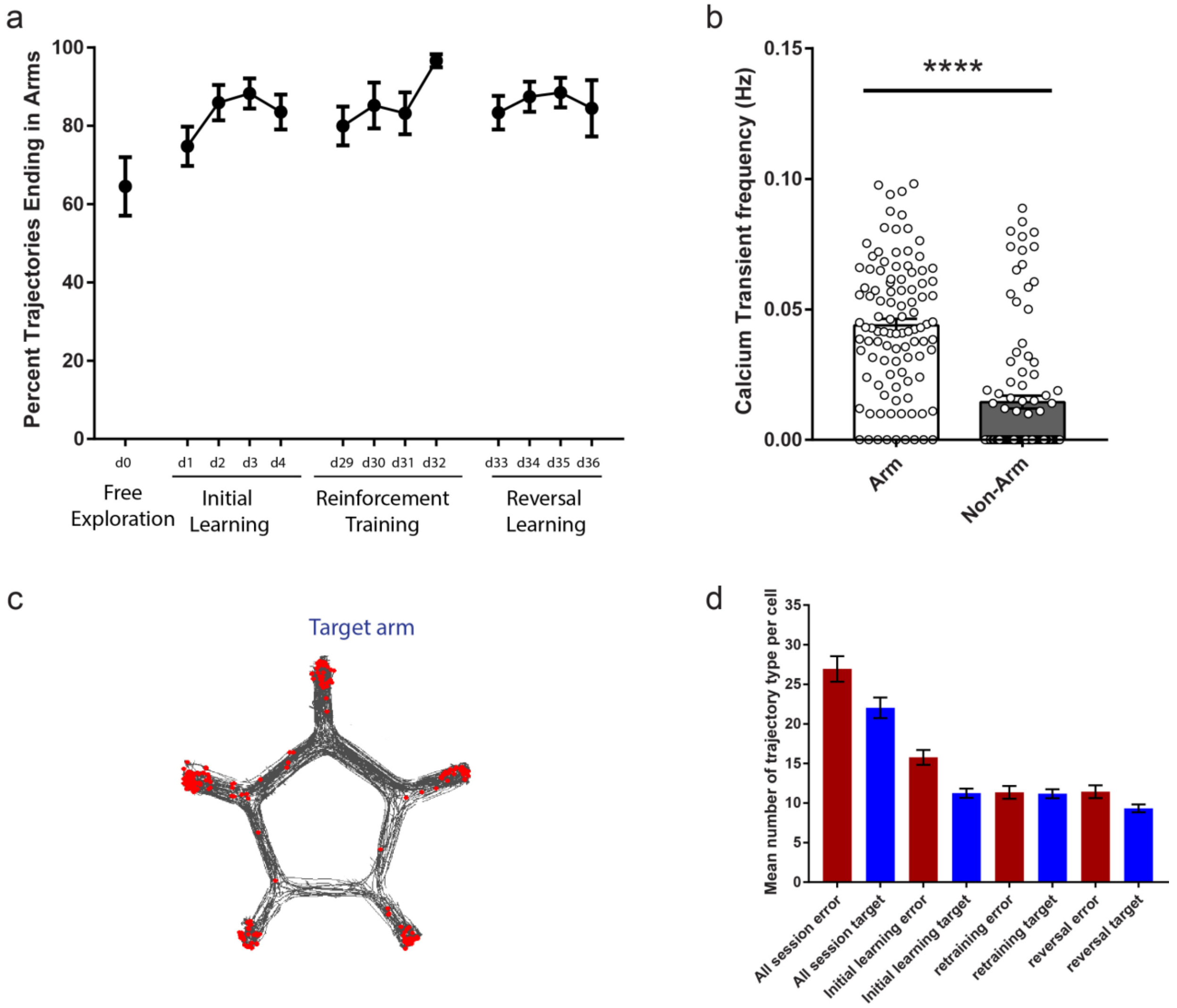


**Supplemental Figure 5 | Trajectory characterization and spatial location of MS glutamate neuron firing.** **a**, Percent of trajectories ending in an arm. In the Dry star maze, 65 to 93% of the trajectories ended in either an error or the target arm. The proportion of arm-directed trajectories increased across learning. **b**, MS glutamate neurons are more active in arms. Mean calcium transient frequency in arms and non-arm region. **c**, Most calcium transients occurs in arms. Example of a mouse path (gray) with superposed calcium transients location (red). d. Mean number of error and target trajectories per cells per testing phase. Error bars represent s.e.m.

**Supplemental Figure 6: *In vitro* characterization of calcium imaging of MS glutamate neurons.**

To verify that changes in calcium signalling reflect changes in the activity of glutamate neurons, *in vitro* recordings were performed in VGLUT2-Cre animals injected with AAV2/9-GCAMP6f-eYFP. Florescent positive neurons were recorded in patch clamp in cell attached mode and the spontaneous activity was recorded both in voltage clamp while the calcium signal was simultaneously captured. Signal was synchronized using a light pulse and combined with a TTL pulse.

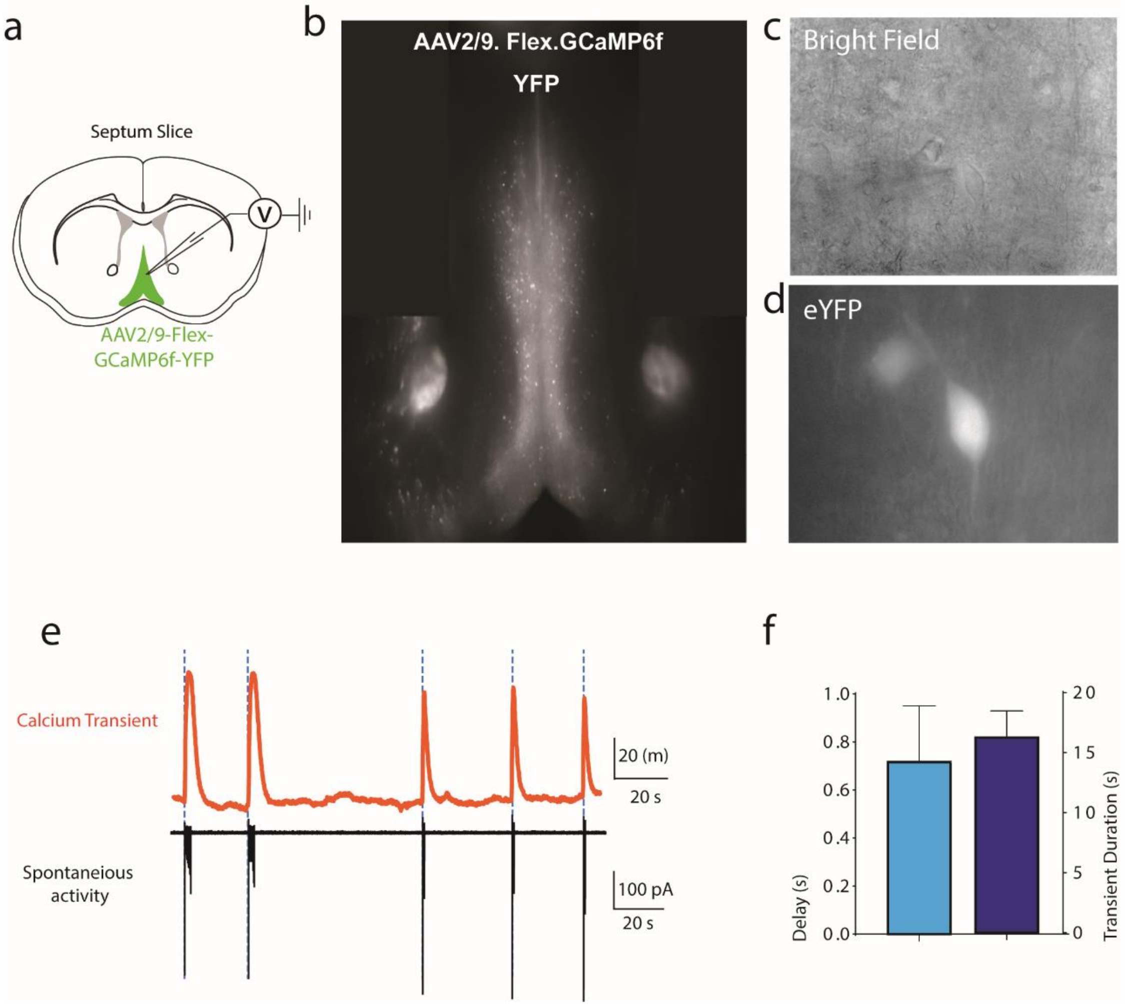


Supplemental Figure 6: (a) Cartoon representation of experimental setup (b) Coronal section showing AAV2/9-GCAMP6f-eYFP expression from MS VGLUT2 neurons. (**c**) Bright field image of target cell. (**d**)

Fluorescence image of target cell. (**e**) Example trace of frequent spontaneous spike bursts that triggers large calcium signals (GCAMP6f) in VGLUT2 neuron recorded in cell-attached mode. (f) Delay between action potential and calcium transient onset and duration of calcium transient (onset mean: 0.7240s ± 0.2265s, duration mean: 16.41±2.062s, *n = 5 cells*).

**Supplemental Figure 7: Specificity of the mouse-line and virus expression**

To ensure that CRE expression is specific to glutamate neurons we compared CRE with VGLUT2 mRNA using florescent in situ hybridization (FISH) and quantified the number of VGLUT2 neurons which are colocalized with CRE and the number of non-specific cells which express CRE mRNA in the absence of VGLUT2 mRNA. In addition, to show the virus distribution across the region of interest, below includes coronal section of the MS ArchT-GFP expression in VGLUT2 neurons.

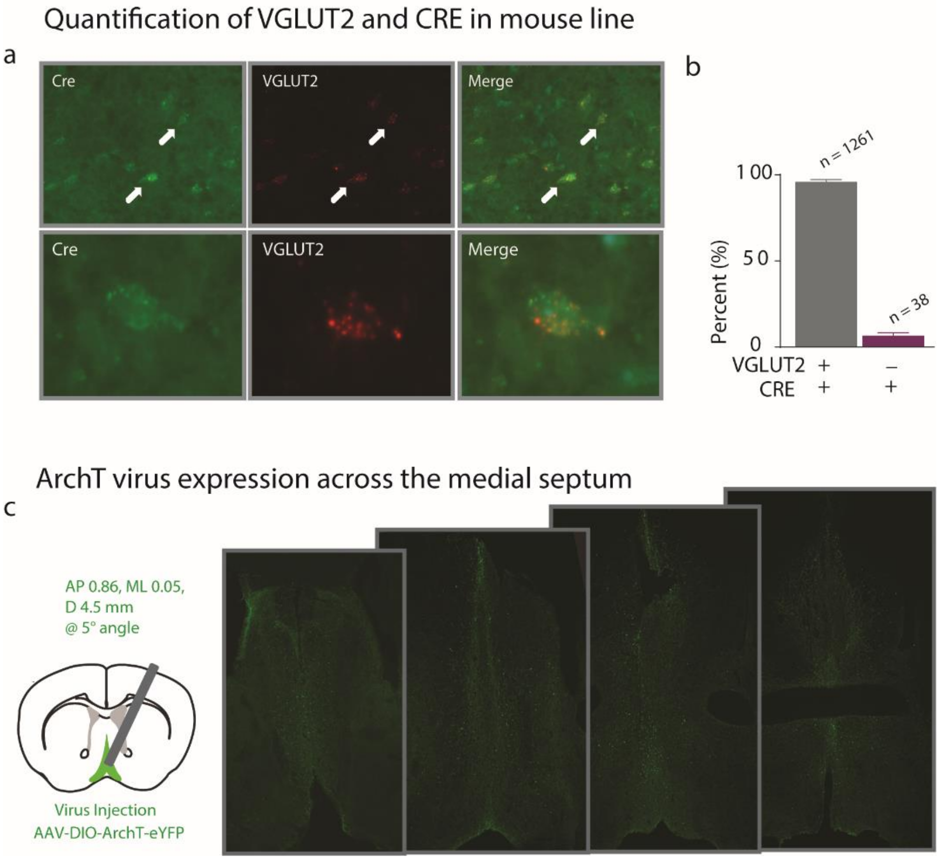


Supplemental Figure 7: (**a**) Cartoon example of experimental setup. (**b**) Percent of neurons which co-express both VGLUT2 and CRE mRNA (mean 95.7 ± 1.607%) and nonspecific expression of CRE mRNA when VGLUT2 is absent (mean 3.431 ± 1.839%). (**c**) Injection coordinates and virus expression across the medial septum.

**_In vitro_ characterization of AAVdj-ArchT in VGLUT2 medial septum neurons.**

To ensure that MS glutamate neurons can be silenced, control experiments were performed *in vitro* to test the opsin. Patch-clamp recordings were performed in VGLUT2-Cre animals injected with AAVdj-ArchT-GFP. Florescent neurons were recorded in whole cell patch-clamp and tested the response to 532 nm light pulses at an intensity of 20 mW.

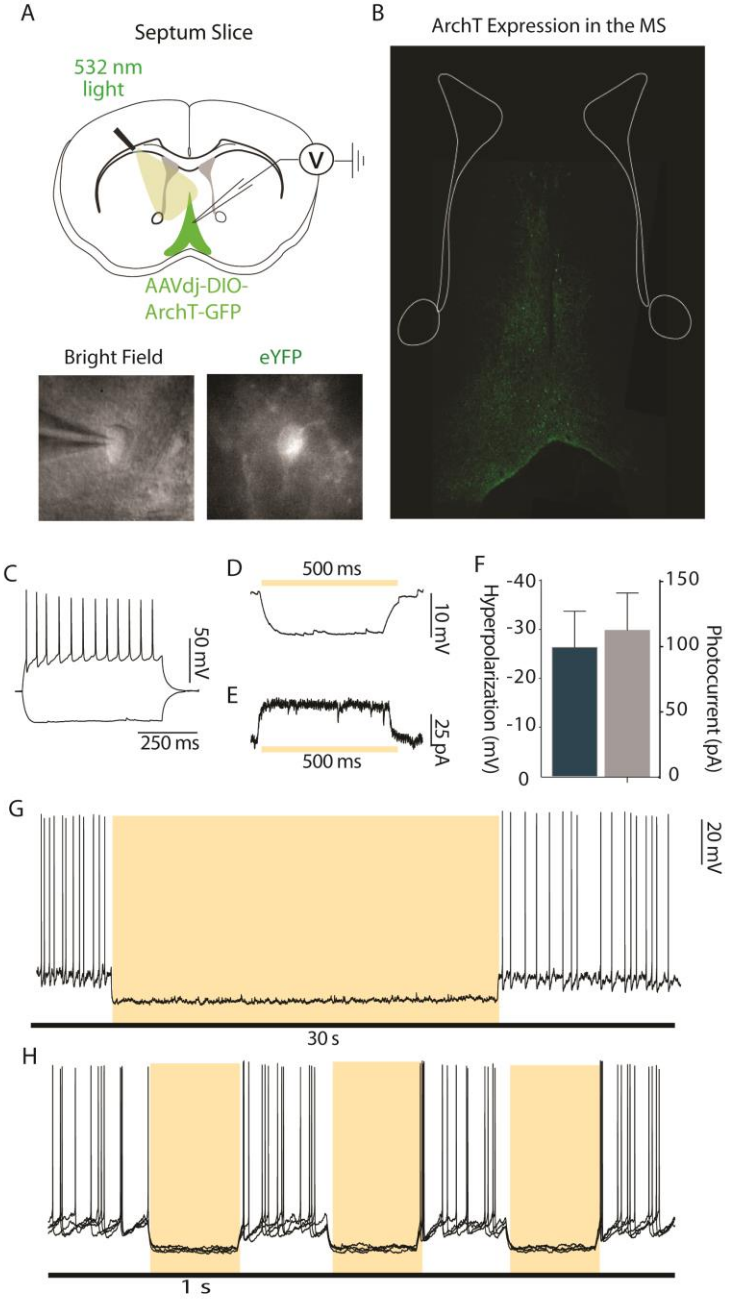


Supplemental Figure 8: (**A.i**) Cartoon example of experimental setup. (**A.ii**) Bright field and fluorescence of neurons patched. (**B**) Virus expression across the medial septum. (**C**) Example of cell characterization. (**D**) Cell hyperpolarization and photocurrent in voltage-clamp (bottom, holding = -70mV) during 500 ms of continuous light stimulation in an ArchT-expressing VGLUT2-expressing (hyperpolarization mean: -26.29 ± 2.07 mV, photocurrent mean: 112.6 ± 28.63 pA, *n* = 7). (**G**) Example trace of 30-second-long continuous light silencing. For both 1 and 30 second, cells were silenced 30 seconds without showing any disruption in firing following the inhibition episode. (**H**) Example trace of 30-second-long continuous light silencing. For both 1 and 30 second, cells were silenced 30 seconds without showing any disruption in firing following the inhibition episode.

**Supplemental Figure 9: Star maze task across 4 day learning task with optogenetic silencing of MS glutamate neurons**

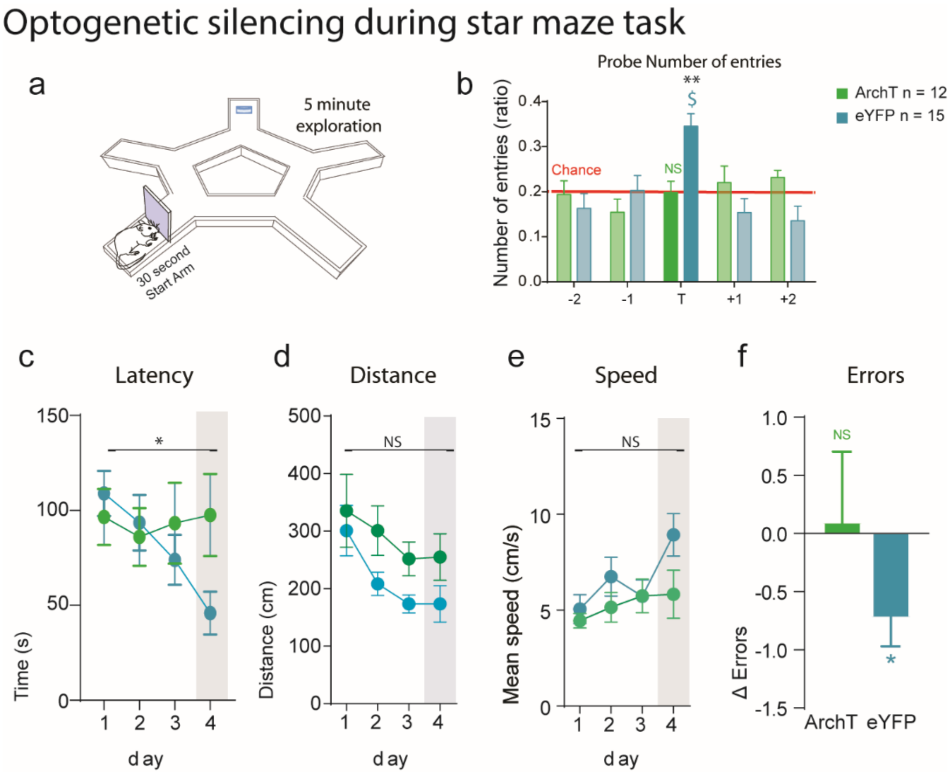


**Supplemental figure 9: (a)** Representation of star maze task. **(b)** Ratio of number of entries in each arm of the maze (experimental group*_Arm interaction_; $F_{(4, 100)} = 4.798$; $P = 0.0014$), ArchT animals had no preference for the target arm compared to chance compared to controls (Holm-Sidak's multiple comparisons). **(c)** Mean latency to target across the 4 day testing period comparing ArchT to control groups (experimental group*_day_; $F_{(3, 74)} = 3.598$; $P = 0.0174$)**. **(d)** Mean distance to target across the 4 day testing period comparing ArchT to control groups (experimental group*_day_; $F_{(3, 74)} = 0.2621$; $P = 0.8525$)**. **(e)** Mean speed to target across the 4 day testing period comparing ArchT to control groups (experimental group*_day_; $F_{(3, 60)} = 1.545$; $P = 0.2121$) **. **(f)** Change in errors to target from day 1. (ArchT mean 0.1042±0.6, one sample t test, t=0.1733, p=0.8656, eYFP -0.73±0.24, one sample t test, t=3.109, p=0.0077). Error bars represent s.e.m. *P<0.05; **P<0.01; ***P<0.001 (_Holm-Sidak's multiple comparison_). $P<0.05; $$P<0.01 (One-sample t-test vs. chance).

**Note** Due to video recording issues, one datapoint was not recorded in the eYFP and so a mixed-effects analysis was performed in place of two-way ANOVA.

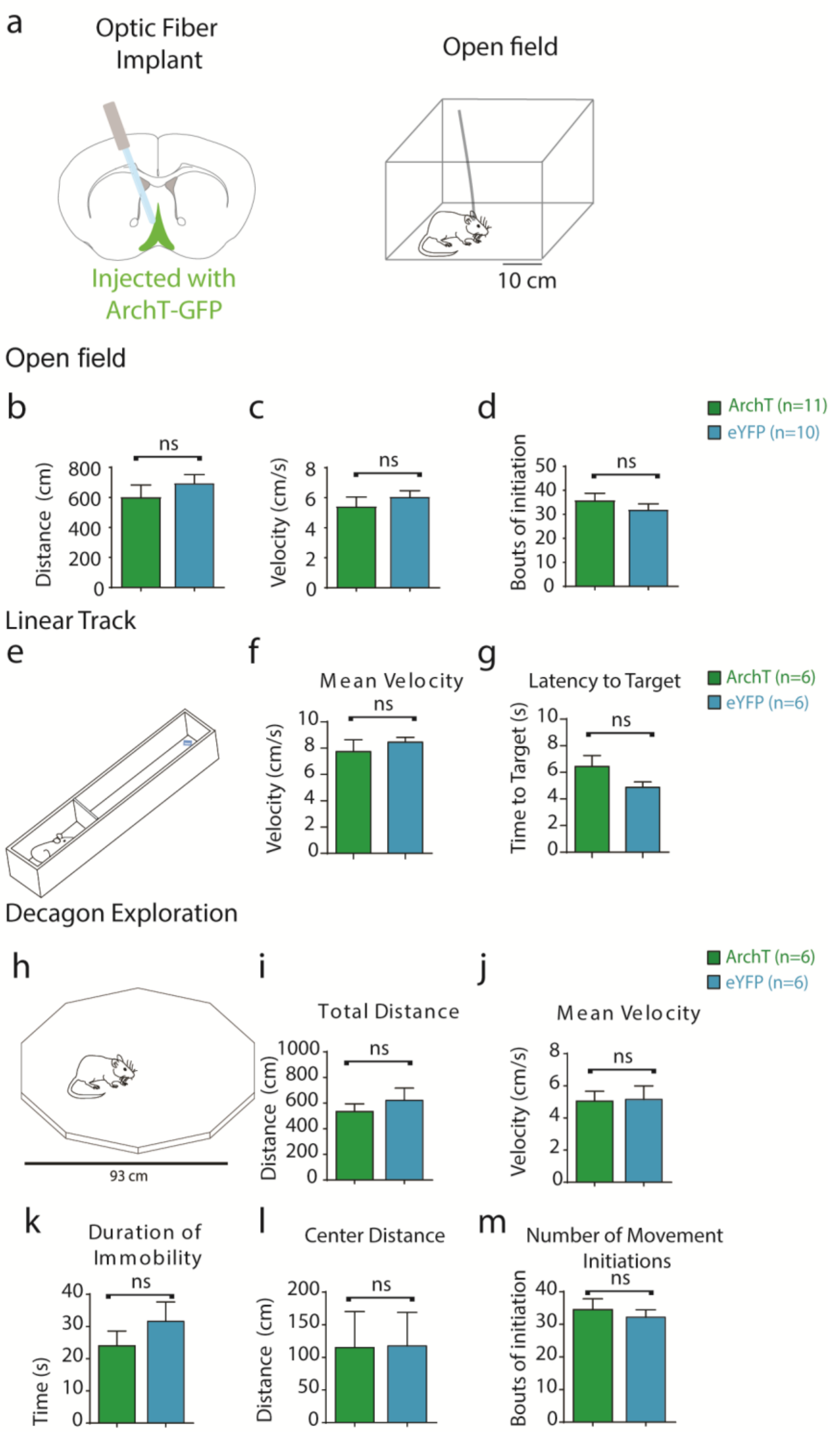


**Supplemental Figure 10: Free exploration during optogenetic silencing**

(**a**) (left) Cartoon representation of experimental setup with optogenetic silencing of the medial septum (right) representation of open field experiment. (**b**) Mean distance explored during open field exploration. (ArchT mean 603±80.3, eYFP 695±57.3 ; $t_{(10, 9)}$=0.9201; p=0.3691). (**c**) Mean velocity explored during open field exploration. (ArchT mean 5.43±0.62, eYFP 6.07±0.41; $t_{(10, 9)}$= 0.8461; p=0.4080). (**d**) Total number of locomotor initiation events during open field exploration (ArchT mean 35.91±2.887, eYFP 32.00±2.394; $t_{(10, 9)}$=1.030; p=0.3158). (**e**) Representation of experiment setup with linear track. (**f**) Mean velocity explored during open field exploration, (ArchT mean 7.74±0.9, eYFP mean 8.45±0.37, $t_{(10, 9)}$=0.7318; p=0.4811). (**g**) Delay to water target (ArchT mean 6.445±0.8134, eYFP mean 4.872±0.4083, $t_{(10, 9)}$=1.729; p=0.1146). (**h**) Representation of experiment setup of decagon open field. (**i**) Total distance explored during 2 minute exploration, (ArchT mean 535±60.02, eYFP mean 622±94.89, $t_{(5,5)}$=0.7772; p=0.4551). (**j**) Mean velocity explored during decagon exploration, (ArchT mean 5.04±0.62, eYFP mean 5.16±0.84, t=0.1084; p=0.9158). (**k**) Duration of immobility during exploration (ArchT mean 23.98±4.84, eYFP mean 31.61±6.0, $t_{(5,5)}$=1.011 p = 0.3360). (**l**) Distance travelled through 50% central distance of open field (ArchT mean 115±55.28, eYFP mean 117±51.37, $t_{(5,5)}$=0.03456; p = 0.9731). (m) Number of movement initations (ArchT mean 34.50±3.324, eYFP mean 32.17±2.272, $t_{(5,5)}$=0.05795; p = 0.5751). unpaired t-test.
